# Intestinal Epithelial Heat Shock Protein 25/27 integrates host and microbial drivers of mucosal restitution following inflammatory injury

**DOI:** 10.1101/2022.06.30.498349

**Authors:** Candace M. Cham, Jeannette S. Messer, Joash Lake, Xiaorong Zhu, Yun Tao, Lei He, Christopher R. Weber, Fanfei Lin, Zhanghan Dai, Jinlu Tong, Sara Temelkova, David T. Rubin, Cambrian Liu, Eugene B. Chang

## Abstract

Mucosal healing following inflammatory injury is poorly understood and often neglected, despite being the best indicator of long-term outcomes in inflammatory bowel diseases. We report here that the enigmatic small molecular weight heat shock protein, Hsp25 (the human form is Hsp27), plays a vital role in converging microbial and host factors to promote pSTAT3-mediated mucosal healing. In wild type mice, the proximal-to-distal gradient of intestinal epithelial cell (IEC) Hsp25 expression is dependent on microbial cues. Patients with left-sided ulcerative colitis, however, show reduced levels of Hsp27 expression in both uninvolved and involved areas compared to normal colons of non-IBD patients. In mice with global or IEC-specific Hsp25 gene-targeted deletion, impaired mucosal healing with development of hallmarks of chronic disease are observed following DSS-induced or TNBS-induced colitis, whereas mucosal restitution is accelerated in IEC-specific overexpressing Hsp25 transgenic mice. In colonic IECs derived from these murine lines, Hsp25 binds and stabilizes a phospho-STAT3/YAP nuclear complex stimulated by IL-22 to sustain its wound healing gene programming. Thus, our findings provide insight into the mechanism of action of IEC Hsp25/27 in integrating host and microbial drivers of mucosal restitution, which can be leveraged to develop novel approaches for achieving and maintaining remission in complex immune disorders like IBD.

## Introduction

Mucosal healing is an essential physiological response to inflamed and injured intestinal mucosa in many diseases, including ischemic colitis, radiation-induced mucosal injury, and inflammatory bowel diseases (IBD)^1^. In fact, successful and complete mucosal healing is associated with better clinical outcomes, longer remission, and lower risk of complications like fibrosis, bleeding, and colorectal cancer^2^. There is also evidence that gut microbes play a major role in this process by limiting intestinal damage and promote wound healing in the intestinal tract^1,3^. Yet, critical gaps in knowledge remain, including if and how microbial and host drivers of mucosal healing work independently or coordinately. In this regard, we have found that a small molecular weight heat shock protein, Hsp25/27 (Hsp27 is the human homolog) expressed by colonic epithelial cells connects host and microbial factors to promote phosphorylated-STAT3 mediated mucosal healing following injury of the gut. In inflammatory bowel diseases, we find that there is decreased expression in both uninvolved and involved mucosa of Crohn’s disease compared to non-IBD normal colonic mucosa, albeit to a great extent in the latter. As we do not observe decreases in Hsp25 expression with increasing inflammation in the Dextran Sodium Sulfate (DSS)-induced murine model of acute, self-limited experimental colitis, it raises the question whether there is an inherent defect or inadequate mucosal Hsp27 expression in human IBD which would impair both host and microbial drivers needed for mucosal healing.

While a myriad of factors are associated with IBD, the gut microbiota is inextricably linked to the pathogenesis of these diseases^4^. Commensal bacteria in the intestinal tract have been shown to reduce chemically induced colitis by promoting intestinal barrier integrity^5^. The gut microbiota influences homeostatic function, regulation of immune cells and regulation of IEC proliferation^3,6,7^. The cross-talk between IECs and the gut microbiota greatly influences the composition of the gut microbiota^8^. In turn, the gut microbiota is known to regulate the induction of anti-inflammatory cells such as Regulatory T Cells and promote the induction and secretion of cytokines such as IL-22^9–11^. Thus, understanding the molecular mediators that enable to IECs cross-talk with the gut microbiota is critical to developing novel therapeutics for promoting mucosal wound healing and ultimately remission in IBD patients.

Heat shock proteins (HSPs) are traditionally thought of as molecular chaperones. They are assumed to promote function and stability with other molecules to confer cytoprotective functions^12–15^. However, novel insights from our studies demonstrates novel roles for HSPs in maintaining intestinal homeostasis^16,17^. Hsp25 and Hsp27 are murine and human homologs, respectively, and are encoded by the *Hspb1* gene, but we denote the gene product as Hsp25/27 for simplicity, as the homologs are nearly identical in amino acid sequence, tertiary structure, and function^18^. In the gut, their constitutive expression in the colonic epithelium is driven by microbial signals. ^12–14^Yet, the physiological role of Hsp25/27 largely remains enigmatic. For example, Hsp25-deficient mice in our studies appear normal and exhibit no visible phenotype. However, we have been able to derive key insights on the role of induced Hsp25 in the Dextran Sodium Sulfate (DSS)-induced colitis using conditional Hsp25 gene-targeted deletion and forced expression mutant mice models. Unlike “classical” induced Hsps like Hsp70 which are cytoprotective and can mitigate the initial stages of mucosal inflammation and injury^12,19^, Hsp25 confers little cytoprotection in our studies. Instead, we noted that there is a significant impairment in mucosal healing during the late “recovery phase” among DSS-treated global and intestinal epithelial cell (IEC)-specific Hsp25 KO mice. In short, Hsp25 KO mice cannot achieve adequate mucosal healing in an acute self-limited colitis model which then goes on to a chronic state. Strikingly, when IEC-specific Hsp25 expression is forced in mice (Hsp25^+IEC^) through a constitutive-CAG promoter driven Hsp25 transgene, mucosal healing is accelerated compared to both WT and Hsp25 KO animals. Our findings further show IEC Hsp25 is a critical integrator of host and microbial drivers of mucosal restitution following inflammation-induced injury.

## Results

### Physiological colonic epithelial Hsp25/27 expression is dependent on microbial signals and exhibits a proximal to distal gradient

The colonic microbiota is essential for the induction and maintenance of colonic epithelial Hsp25, as shown in immunostaining images of Figure 1. SPF mice which show predominantly Hsp25 expression in surface epithelium of proximal colon (Figure 1A) and less in distal SPF colonic mucosal (Figure S1) which likely stems from the very diverse gut microbiota in the proximal colon that include flagellated and fermentative gut microbes^20,21^. In contrast, germ-free (GF) mice do not exhibit colonic Hsp25 expression (Figures 1A and S1, bottom panel).

**Figure 1.**
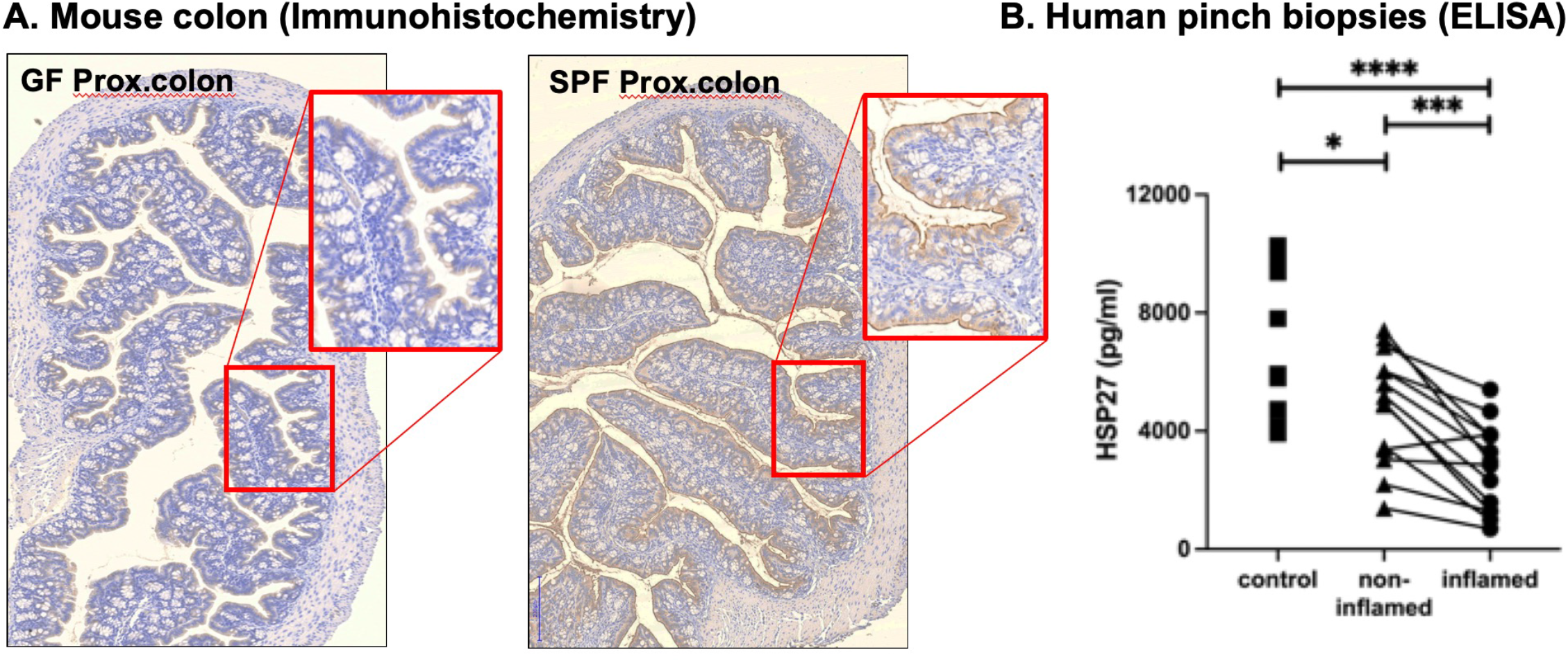
Colonic Hsp25/27 expression requires microbial signals and is decreased in inflamed mucosa of IBD patients. (A) Hsp25 expression is greater in proximal colon IECs from SPF mice versus GF mice (note brown staining of IECs of SPF inset). (B) Hsp27 protein expression (by ELISA) is significantly decreased in inflamed left-sided ulcerative colitis mucosa compared to paired, adjacent non-inflamed regions and non-IBD control tissues.

Human colonic mucosa expression of Hsp27 in mucosal pinch biopsies from non-IBD controls and IBD subjects was also measured by ELISA (Figure 1B). Notable in IBD subjects is that Hsp27 expression is significantly less in matched involved and non-involved areas compared to non-IBD controls and in inflamed regions vs non-inflamed regions of individual patients (connecting lines indicating the same patient). These differences likely arise from differences in regional microbiota and their products associated with IBD gut dysbiosis, but the possibility that IBD patients have an inherently compromised Hsp27 response cannot be ruled out.

### IEC Hsp25 is essential for mucosal healing in the DSS-colitis model

We employed conditional genetic Hsp25 transgenic and KO mice established by our laboratory (Table 1) to derive key insights on the role of induced Hsp25 in human IBD and experimental mouse models of colitis. In Figure 2A, mucosal Hsp25 protein expression increases significantly as the inflammatory phase of DSS colitis transitions to the recovery phase (∼Day 8). Unlike “classical” induced Hsps like Hsp70, which are cytoprotective and can mitigate the first stages of mucosal inflammation and injury^22^, Hsp25/27 appears to have minimal effects in this initial inflammatory phase. Slight, but significant differences were observed in clinical disease activity index (See Figure 2B) between DSS-treated Hsp25 KO and wild type mice, greater among male mice. However, by more objective criteria of histologic and LCN2 indices, no significant differences among these groups were observed during the inflammatory period. In contrast, a significant impairment in outcomes during the late “recovery phase” (Days 8+, green shaded area) among DSS-treated Hsp25 KO mice (Figure 2C) is observed accompanied by histological hallmarks of chronic colitis such as mucosal crypt branching, atrophy, and erosions (at Days 30 and 45, well after healing was achieved in wildtype animals where nearly complete recovery is observed by day 19). In short, in absence of Hsp25, acute self-limiting DSS-induce colitis in WT mice becomes chronic.

**Table 1:** WT and Hsp25 gene modified mouse lines (all C57Bl/6) and TARGATT vector to force IEC-specific Hsp25 gene expression.

| Symbol | HSP25 Phenotype | Genotype | Description |
| --- | --- | --- | --- |
| Hsp25 WT | Wildtype | Hsp25<sup>fl/fl</sup> | Normal expression of Hsp25<br>LoxP sites flanking exons 2 and 3 of Hsp25 gene<br>Parental line of some mouse lines listed in this table |
| Hsp25 KO | Global knockout | Hsp25<sup>-/-</sup> | Deleted *hspb1* gene encoding Hsp25 in all tissues<br>Generated by crossing Hsp25<sup>fl/fl</sup> mice with (sperm-specific) Sycp1-Cre transgenic mice, thus creating progeny whose germline lacks Hsp25; Sycp1-Cre transgene is bred out |
| Hsp25<sup>+IEC</sup> | Conditional overexpression of Hsp25 in the intestine | Hsp25<sup>TARGATT</sup> x Vil1::Cre (JAX strain #020282) | Constitutive overexpression of IEC Hsp25<br>Hsp25<sup>TARGATT</sup> transgenic: LoxP sites flanking STOP cassette inserted upstream of Hsp25 gene plasmid (TARGATT) |
| Hsp25<sup>ΔIEC</sup> | Conditional deletion of Hsp25 in the intestine | Hsp25<sup>fl/fl</sup> x Vil1::Cre | Deletion of HSP25 in the intestinal epithelial cells<br>Generated by crossing Hsp25<sup>fl/fl</sup> mice with Villin1-Cre transgenic mice |

|  |  |  |  |
| --- | --- | --- | --- |
| Symbol | HSP25 Phenotype | Genotype | Description |
| Hsp25 WT | Wildtype | Hsp25<sup>fl/fl</sup> | Normal expression of Hsp25<br>LoxP sites flanking exons 2 and 3 of Hsp25 gene<br>Parental line of some mouse lines listed in this table |
| Hsp25 KO | Global knockout | Hsp25<sup>-/-</sup> | Deleted *hspb1* gene encoding Hsp25 in all tissues<br>Generated by crossing Hsp25<sup>fl/fl</sup> mice with (sperm-specific) Sycp1-Cre transgenic mice, thus creating progeny whose germline lacks Hsp25; Sycp1-Cre transgene is bred out |
| Hsp25<sup>+IEC</sup> | Conditional overexpression of Hsp25 in the intestine | Hsp25<sup>TARGATT</sup> x Vil1::Cre (JAX strain #020282) | Constitutive overexpression of IEC Hsp25<br>Hsp25<sup>TARGATT</sup> transgenic: LoxP sites flanking STOP cassette inserted upstream of Hsp25 gene plasmid (TARGATT) |
| Hsp25<sup>ΔIEC</sup> | Conditional deletion of Hsp25 in the intestine | Hsp25<sup>fl/fl</sup> x Vil1::Cre | Deletion of HSP25 in the intestinal epithelial cells<br>Generated by crossing Hsp25<sup>fl/fl</sup> mice with Villin1-Cre transgenic mice |

**Figure 2.**
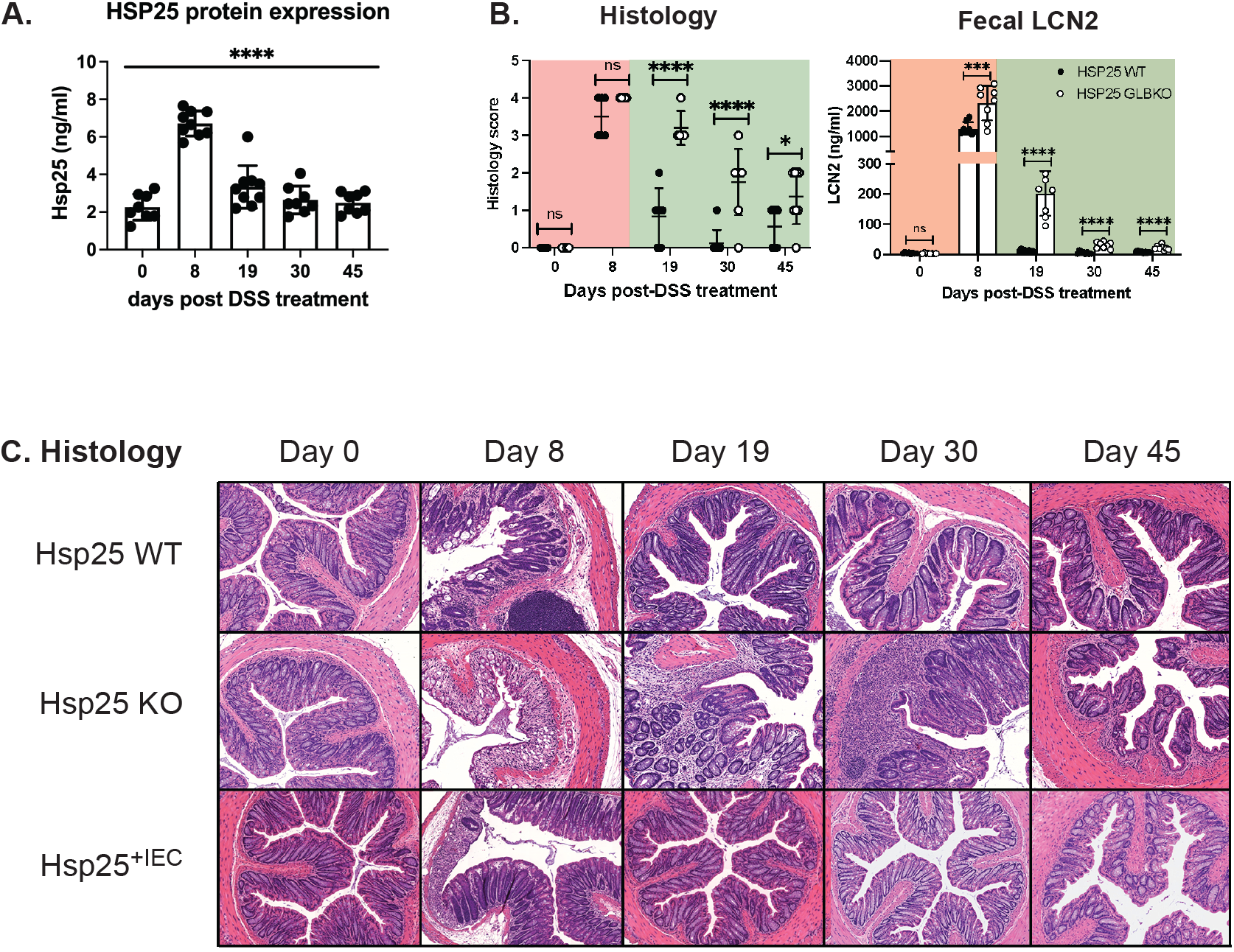
Hsp25 is essential for mucosal healing from DSS-induced colitis. (A) Hsp25 expression is significantly increased in both inflammatory and recovery phases of DSS colitis in WT mice; (B) Histology scores and LCN-2 levels for Hsp25 WT and KO mice treated with DSS during the inflammatory (red) and recovery (green) phases (*p,0.05, ****p<0.0005); (C) Colonic mucosal histology at different time points of DSS inflammatory and recovery phases.

Strikingly, IL-22 levels are greater in Hsp25 KO mice compared to WT, suggesting a post-IL-22 receptor defect rather than a lack of IL-22 production in Hsp25 KO mice (Figure S3A). When IEC-specific Hsp25 expression is forced in mice (Hsp25^+IEC^) through a constitutive, epithelial-specific CAG promoter-driven Hsp25 transgene (see Figure 2C, bottom row), Hsp25^+IEC^ mice exhibit accelerated mucosal healing (and decreased injury) compared to both WT and Hsp25 KO animals (see Day 8 of Figure 2C). Gut epithelial Hsp25 expression is sufficient for this response, as compromised mucosal healing and persistence of DSS-induced colitis is observed at D19 in Hsp^ΔIEC^ mice where targeted deletion of intestinal epithelial specific Hsp25 was performed (Figures 2A, S2A). Further supporting the compromise in mucosal healing is Ki67 staining which shows significant impaired proliferative response in Hsp25 KO mice (Figure S3B). Similar findings of compromised mucosal healing are observed in the TNBS-induced model of colitis where a wound reparative response can be seen in Hsp25^f/f^ mice (Figure 3B) that is associated with an increased IL-22 mucosal response in comparison with baseline expression of Hsp25^f/f^ mice not treated with TNBS (Figure 3C). In contrast, TNBS-treated Hsp25^ΔIEC^ counterparts exhibited a significantly increased IL-22 mucosal levels and a lack of histologic wound healing response (Figures 3C,D) compared to Hsp25^f/f^ mice. As will be shown below, this paradoxical increase in IL-22 mucosal levels in Hsp25^ΔIEC^ mice appear to arise from a post-receptor block due to impairment of IEC pSTAT3 signaling.

**Figure 3.**
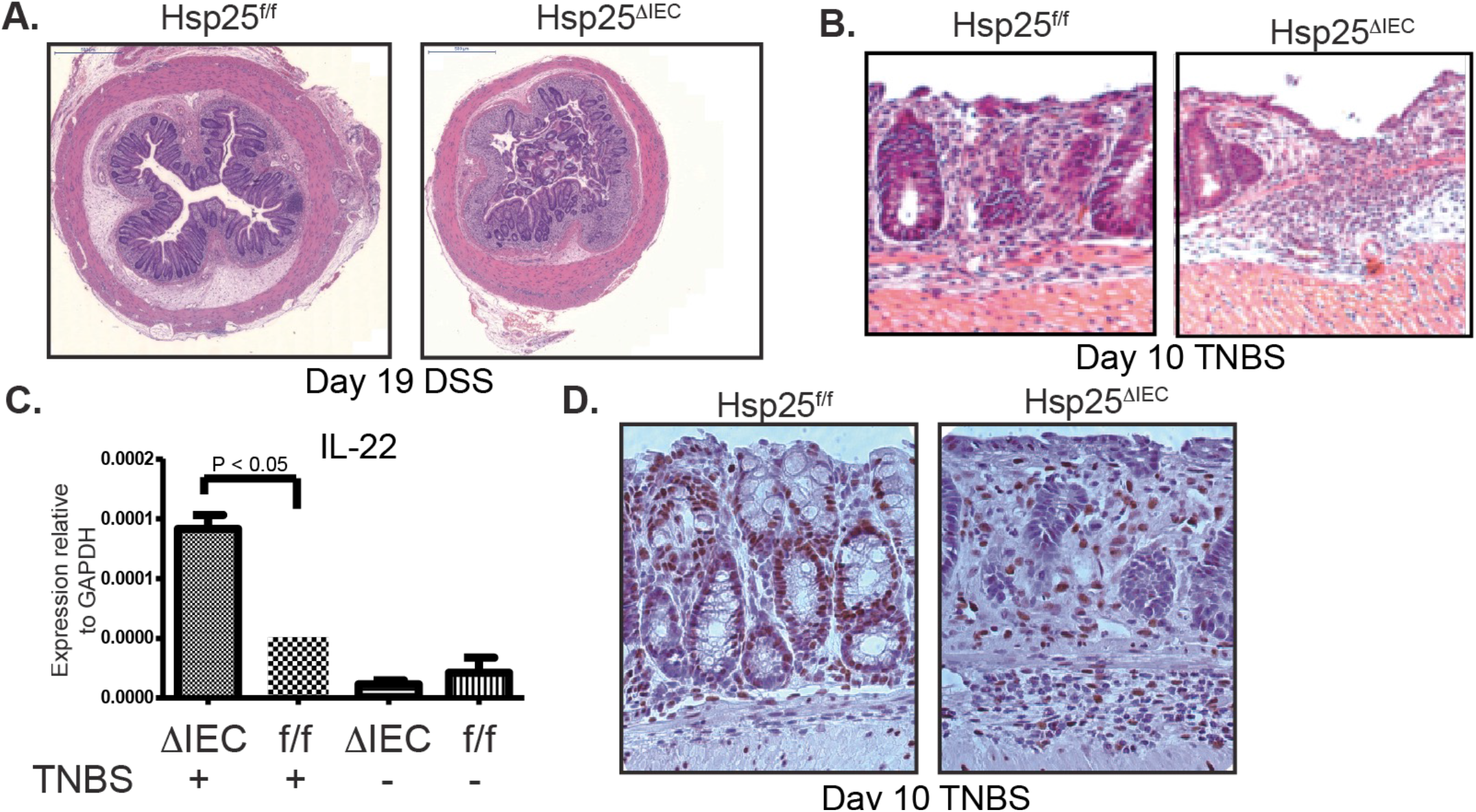
Hsp25^ΔIEC^ exhibit impaired mucosal healing responses to TNBS- and DSS-induced colitis compared to Hsp25^f/f^ controls. H&E of distal colon from Day 19 DSS (A) and Day 10 TNBS (B). qRT-PCR for distal colonic mucosal IL-22 on Day 10 TNBS (C). IHC for pSTAT3 Y705 (brown IEC nuclei staining) in colonic mucosa of Hsp25^f/f^ vs. Hsp25^ΔIEC^ mice at D10 TNBS treatment (D).

### Hsp25 associates with phosphorylated Y705 STAT3 (pSTAT3), a transcriptional regulatory factor involved in inflammation and mucosal healing

We next examined the molecular mechanisms through which Hsp25/27 potentially mediates epithelial responses to colitis and mucosal healing. We observed reduced pSTAT3 Y705 in mucosal scrapings of DSS-treated Hsp25 KO mice compared to WT mice at D19 post-DSS (Figure 4A). Several studies have shown that Hsp25 interacts with pSTAT3^14,23,24^. We confirm this and further show that IL-22 induced phosphorylation of STAT3 is at both Y705 and S727 of colonic epithelial organoids of Hsp25 WT mice, whereas it is diminished in the absence of Hsp25 (shown only for Y705 in Figure 4). Total STAT3 expression, however, is no different between the two groups. Figure 4B shows that both IL-22-stimulated nuclear and cytosolic pSTAT3 of colonic organoids are decreased in absence of Hsp25 (Nup98 is a nuclear marker). We further demonstrate that Hsp25 and pSTAT3 are co-immunoprecipitated using a Western blot assay, indicative of their interaction and binding in WT Hsp25 IECs following IL-22 stimulation (Figure 4C).

**Figure 4.**
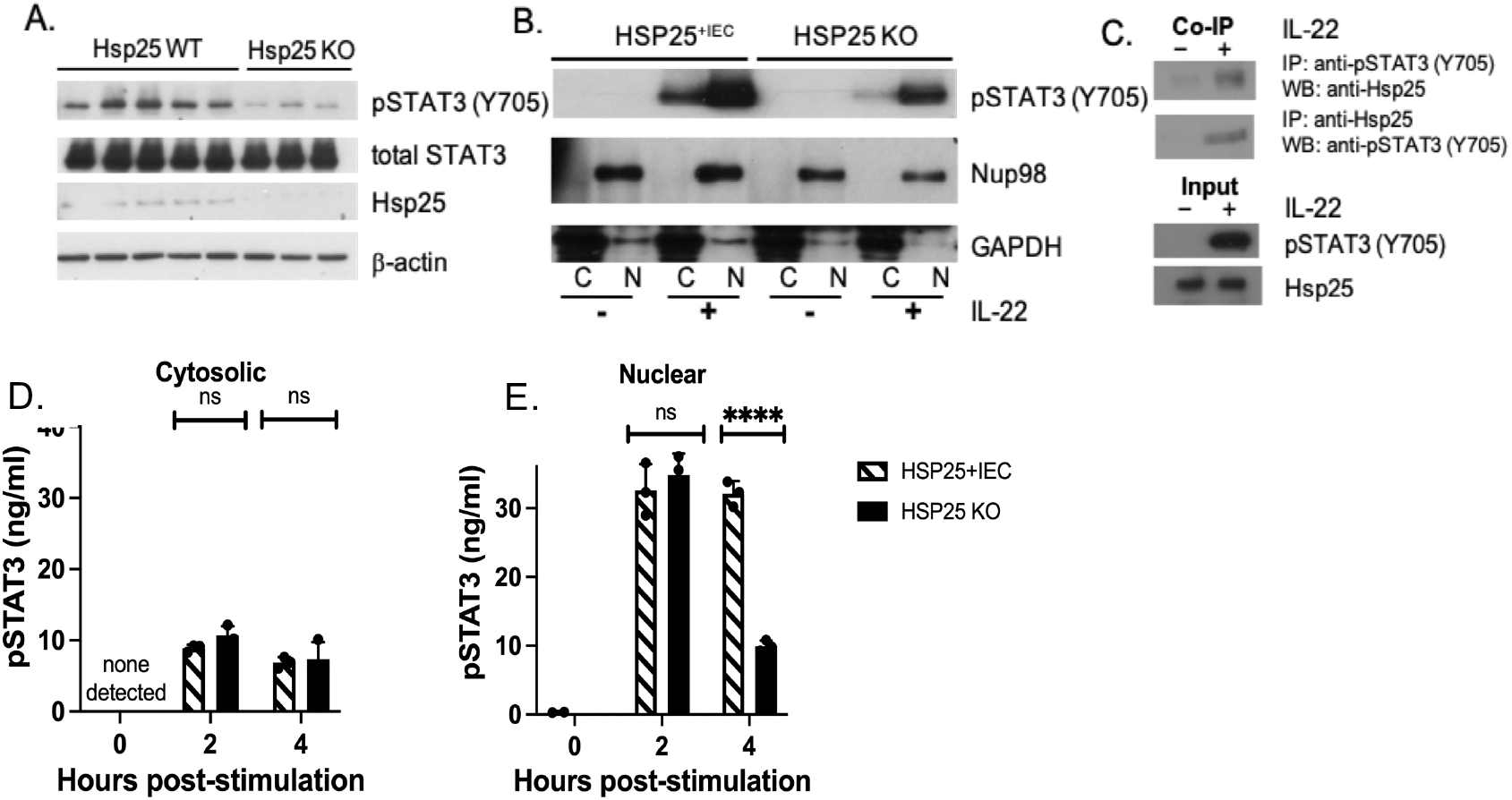
Hsp25 associates with nuclear phosphorylated STAT3 (pSTAT3 (Y705)), a transcriptional regulatory factor involved in both inflammation and mucosal healing. (A) pSTAT3 (Y705) is reduced in D19 mucosal scrapings of DSS-treated Hsp25 WT and KO mouse colons. (B) After 2 hours of IL-22 stimulation (30 ug/ml), Hsp25 KO colon IECs exhibit reduced pSTAT3 (Y705) compared to Hsp25^+IEC^ colon IECs. (C) In IL-22 stimulated IECs isolated from Hsp25^+IEC^ colons, Hsp25 associates with pSTAT3 Y705. (D and E) Cytosolic and nuclear fractions from IL-22 stimulated Hsp25^+IEC^ and Hsp25 KO colonic IECs were measured by ELISA.

When a time course study is performed (Figure 4D) for IL-22 stimulated cytosolic and nuclear pSTAT3 expression at 2 hours are no different between Hsp25^+IEC^ and Hsp25 KO colonic organoids, but in absence of Hsp25, pSTAT3 levels are significantly decreased by 4 hrs only in Hsp25 KO cells, suggesting Hsp25 is needed to stabilize and sustain the nuclear pSTAT3 response, but is not involved in the initial phosphorylation event (Figure 4D, right panel). While the decreased pSTAT3 at 4h could be due to degradation in absence of Hsp25, the finding that total cellular STAT3 is unchanged (Figure 4A), suggesting that Hsp25, as a molecular chaperone, protects against phosphatase activity. In both colonic scrapings of D19 DSS-treated Hsp25 WT mice and in IECs isolated from Hsp25^+IEC^ mice, phosphorylation of STAT3 is observed at Y705 (Figure 4A,B), but, in the absence of Hsp25, phosphorylation at both sites is reduced. As total STAT3 is unchanged, we again posit that more rapid dephosphorylation of pSTAT3 rather than its protein degradation is involved in absence of Hsp25, resulting in impairment of mucosal restitution.

### Hsp25/27 as a determinant of nuclear pSTAT3 binding with reparative transcriptional activator, YAP

Two highly related transcriptional activators, YAP and TAZ, are important mediators of intestinal mucosal wound healing through their integration of cell polarity and mechanical cues from growth factor signaling and inflammation^25^. In vitro, YAP/TAZ has been shown to dedifferentiate committed cells back to a progenitor and stem cell state^26^ and reprogram colonic epithelium cells by linking ECM remodeling to tissue regeneration^27^. As shown in Figure 5A, the co-immunoprecipitation of YAP with pSTAT3 Y705 is significantly decreased in Hsp25-deficient mouse colonic IECs. This finding provides a plausible explanation for how pSTAT3 activation by reparative cytokines is compromised in inflammatory conditions where Hsp25/27 is absent or decreased.

**Figure 5.**
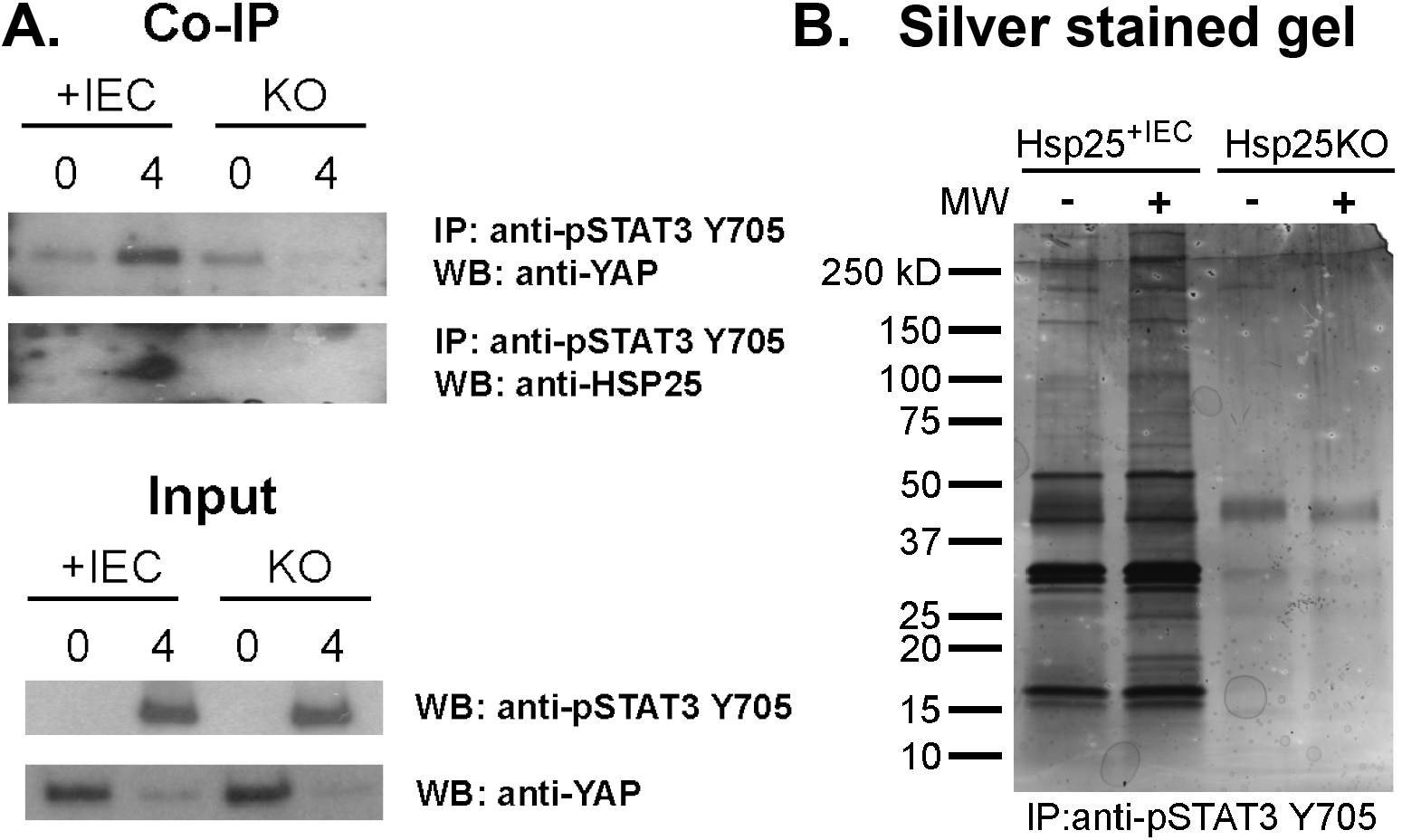
YAP is in a transcriptional complex with pSTAT3 Y705 and Hsp25. (A) Co-IP of Hsp25^+IEC^ and Hsp25 KO colonic IECs stimulated for 4 hours with IL-22 (30 ug/ml). (B) Representative polyacrylamide gel of proteins that immunoprecipitated with anti-pSTAT3 Y705 antibody following IL-22 stimulation for 3 hours, as detected by silver stain.

## Discussion

Mucosal healing, particularly in IBD, is not well understood – but is an important aspect for therapeutics of immune and inflammatory diseases that has received little attention relative to immunosuppressive and immunobiologic interventions. Currently, complete endoscopic healing is the end point that defines clinical remission, but we have no measures to promote mucosal restitution, particularly in inflammatory bowel diseases. Here, we report a novel role for the small molecular weight heat shock protein, Hsp25/27, whose roles and functions are incompletely understood. Several prior studies have suggested that Hsp25/27 is involved in tumor progression and metastasis^28,29^, protection of cells against apoptosis under stressful conditions^30,31^ and smooth muscle development and contractility^24,32–34^, but many of these studies predate the development of genetic experimental models and based conclusions on observational data. Using both *in vitro* and *in vivo* approaches, the latter involving cell specific Hsp25 gene-targeted deletion and overexpression murine models, we report that Hsp25/27 is an essential facilitator of the transcription factor STAT3 (Signal transducer and activator of transcription 3) which functions to converge many cooperative gut microbial and host signals that promote intestinal mucosal restitution following immune and inflammatory stress and injury. STAT3 has been shown to control cell growth, division, migration, and apoptosis to pro-inflammatory actions and its phosphorylation and activation by IL-22 promotes mucosal healing^35–41^. Conversely, STAT3 is involved in pro-inflammatory cytokine signaling through the Janus kinase/signal transducer and activator of transcription (JAK/STAT) signaling pathway that is associated with autoimmune disorders and various cancers^42^. In this regard, inhibitors of the JAK-STAT pathway are currently being used for these diseases, primarily to counter the activated immune states.

Physiological expression of Hsp25 in murine colonic epithelium is dependent on cues from the gut microbiota such as short chain fatty acids (particularly butyrate) and flagellin (through TLR5 activation)^16,17^, as levels are undetectable in germ-free mice (Figure 1A). Presumably, regional differences in microbiota and their products account for the proximal to distal gradient of Hsp25 expression of Hsp25^21^. In human IBD (ulcerative colitis), however, we find that mucosal Hsp27 expression is significantly decreased in both non-involved and involved areas of patients (greater in the latter) with left sided ulcerative colitis compared to non-IBD controls (Figure 1B). We attribute these differences to changes in regional colonic microbiota and products like SCFAs that are needed to sustain levels of Hsp27, but we cannot exclude the possibility that IBD patients have inherently decreased Hsp27 expression. DSS-treated mice, for example, exhibit increased Hsp25 expression in the height of mucosal inflammation (D8) and throughout the recovery period (post D8) (Figure 2A).

Our study highlights a unique and previously unknown role for Hsp25/27 in mucosal healing following inflammation that integrates microbial and host drivers of mucosal restitution at a critical point of convergence (pSTAT). We show, for instance, that gene-targeted deletion of Hsp25 has minimal effects on colitis development following DSS treatment but is essential for the wound healing response that occurs after Day 8. In fact, mucosal healing is accelerated in IEC-specific Hsp25-overexpressing transgenic mice (Figure 2). With both global and IEC-specific gene-targeted deletion of Hsp25, the main differences with WT and Hsp^+IEC^ transgenic mice are impaired mucosal healing in the recovery phase where there are features of chronic colitis, including mucosal atrophy, decreased proliferation, and branching crypts, immune cell infiltration, and crypt abscesses even as late as 45 days post-DSS colitis induction. In this regard, it is notable that IEC-specific expression of Hsp25 is sufficient by itself for mucosal healing, as recovery is impaired in the Hsp^ΔIEC^ mice but promoted in Hsp^+IEC^ transgenic mice where Hsp25 is constitutively driven even during inflammation.

We also define the mechanisms and targets by which Hsp25/27 integrates host and microbial drivers of intestinal mucosal restitution. First, physiological expression of colonic epithelial Hsp25 expression requires microbial cues from regional taxa that include short chain fatty acids^16^ and TLR-5 activating flagellin^17^. The proximal to distal gradient of colonic epithelial Hsp25 expression is mostly like due to the regional diversity which is more abundant and diverse proximally and where fermentation is greatest^21,43^. Many of these taxa and fermentative processes are significantly decreased in the dysbiosis associated with human IBD, accounting for the decreased Hsp27 expression associated with involved and non-involved regions of patients with left sided ulcerative colitis. However, the possibility that there is an inherent defect in Hsp27 response in IBD subjects cannot be ruled out. Decreased or absent Hsp25/27 expression during the recovery period impacts the actions of host reparative factors such as IL-22 by influencing the sustainability of nuclear phospho-STAT3 stability and possibly direct the selection of nuclear transcriptional complex partners such as YAP and TAZ, both well-recognized transcription factors that associate with pSTAT3 to promote cellular proliferation, survival, and differentiation pathways essential for wound repair^44,45^.

As shown in figure 6, we hypothesize that Hsp25 is critically involved the induction of the wound-repair cytokine IL-22 response, but not the proinflammatory IL-6 response in the colonic epithelium. Upon IL-22 receptor engagement and STAT3 phosphorylation, Hsp25 serves to stabilize pSTAT3, either protecting it from degradation or from dephosphorylation, thereby allowing pSTAT3 to interact with the transcriptional activators YAP/TAZ and intestinal restitution. On the other hand, IL-6 receptor engagement also results in STAT3 phosphorylation, yet signal transduction from the IL-6 receptor is not dependent of Hsp25, and a different set of genes are transcribed. In the absence of Hsp25, pSTAT3 is not sustained, therefore preventing the binding of YAP/TAZ, and resulting in chronic colitis due to the lack of wound healing.

**Figure 6.**
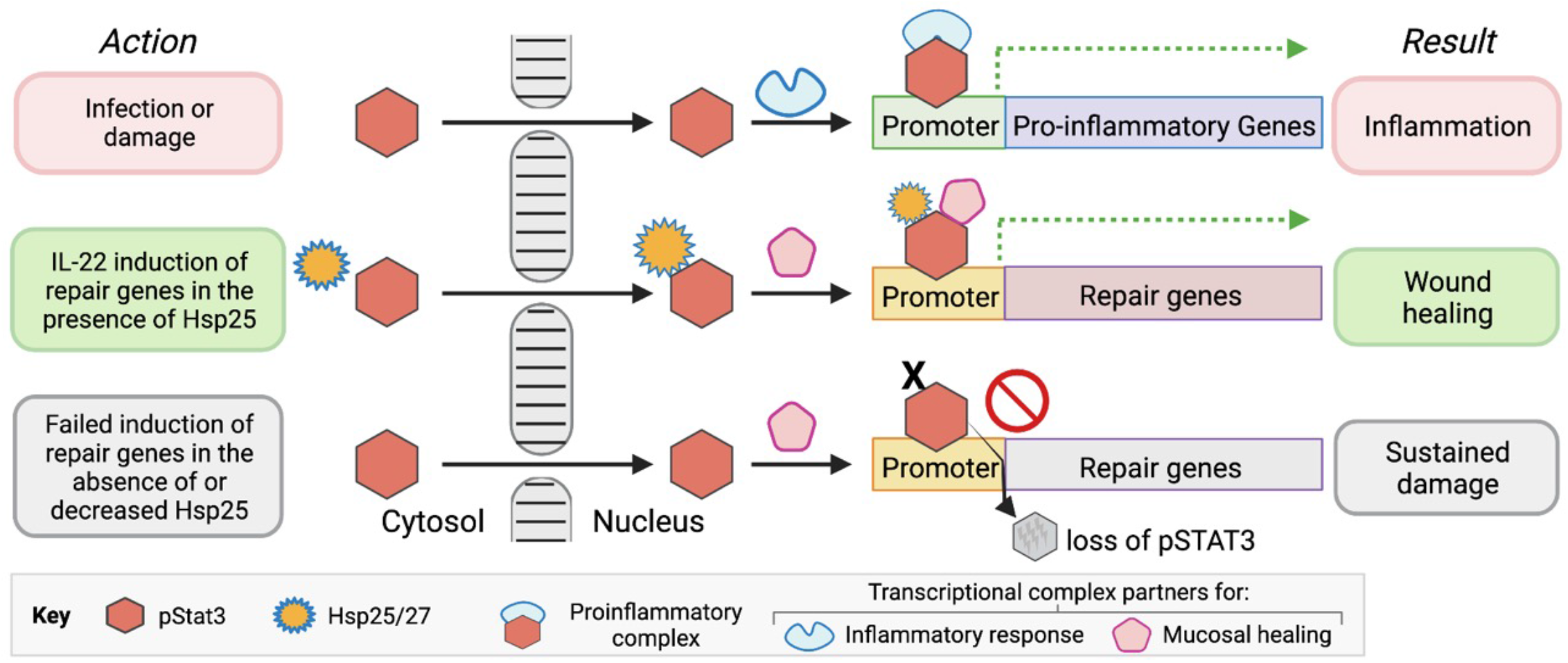
Ligand-stimulated phosphorylation of STAT3 (pSTAT3) can induce a pro-inflammatory or wound healing response depending on the sites of phosphorylation (e.g., S727 vs. Y705) and what protein partners nuclear pSTAT3 assembles with to form transcriptional regulatory complexes that target different gene promoters (top and middle paths). We hypothesize that Hsp25/27 binding with pSTAT3 is dependent on Y705 phosphorylation, which stabilizes it. In addition, it may also influence association with other transcriptional complex partners that activate specific genes essential for mucosal healing (e.g., YAP). In contrast, pro-inflammatory IL-6 appears to preferentially phosphorylate STAT3 S727, which may not promote binding and direction by Hsp25.

In summary, we show an essential role of IEC Hsp25/27 in the convergence of host and microbial signals that drive host mucosal restitution following intestinal inflammation and injury. We also show that the physiological regulation of Hsp25/27 is an important component of this action and that decreased or absent Hsp25 can negatively impact proper and complete wound healing response. While having impairment of the Hsp25/27 expression and response is unlikely to be specific to human IBD, it is likely an important contributor and mediator of IBD chronicity and inability for many patients to achieve clinical remission. In this regard, mucosal Hsp27 gene and protein expression could be useful as predictive markers of response to therapy. Finally, the finding that mucosal restitution in DSS-induced colitis is accelerated in Hsp^+IEC^ transgenic mice suggests that strategies promoting Hsp25/27 gene expression and function represent opportunities for further development.

## Methods

### Mice

All animal protocols and experimental procedures were approved by the University of Chicago Institutional Animal Care and Use Committee (IACUC). A mixture of males and females mice ranging in age from 8 weeks to 30 weeks old were used for all *in vitro* and *in vivo* experiments. Specific pathogen-free (SPF) Hsp25 WT, Hsp25 KO, Hsp25^+IEC^, Hsp25^f/f^, Hsp25^_IEC^ mice were all bred in the University of Chicago animal vivarium. Germ-free (GF) mice were maintained in plastic flexible film isolators (CBC Ltd. Madison, WI, USA). All mice were held under standard 12:12 light/dark conditions.

The genotype of Hsp25 WT mice is Hsp25^f/f^. These mice were generated by the insertion of LoxP sites flanking exons 2 and 3 of Hsp25 (*Hspb1*) gene. To generate mice deficient in Hsp25 in all tissues, Hsp25^f/f^ mice were bred to the sperm-specific Sycp1-Cre transgenic mice, thus creating progeny whose germline lacks *Hspb1*. The Sycp1-Cre transgene is then bred out.

Hsp25^+IEC^ mice were generated by first creating the Hsp25^TARGATT^ transgenic construct: LoxP sites flanking a STOP cassette was inserted upstream of the *Hspb1*-containing plasmid of the TARGATT vector (Applied Stem Cell, Milpitas, CA). This construct was inserted into the Rosa26 locus. Next, Hsp25^+IEC^ mice were crossed with Villin-Cre transgenic mice, resulting in the creation of the Hsp25^+IEC^ line where Hsp25 is overexpressed in the intestine.

Hsp25^_IEC^ mice were generated by crossing Hsp25^f/f^ mice with Villin-Cre transgenic mice. Cre expression is driven by the Villin (intestinal-specific) promoter. Therefore, these mice are conditional knockouts where Hsp25 is expressed normally throughout the body except for the intestine, where there is no Hsp25 expression.

### Human subjects

The study was approved by the Institutional Review Board at the University of Chicago. Subjects were consented for the study at the time of endoscopic examination. For IBD patients, inclusion criteria included subjects aged ≥18 y with a history of ulcerative colitis or Crohn’s disease confirmed by histologic evaluation and a disease extent >20 cm proximal to the anal verge. Healthy subjects were recruited at the time of routine surveillance colonoscopy.

### DSS treatment of mice

Prior to DSS treatment, cage bedding among the experimental mice were mixed once per week for 4 weeks to normalize the microbiomes across genotypes (Miyoshi et al 2019 PeerJ). Mice between the ages of 8-12 weeks of age were given 3% DSS (w/v) in the drinking water for 5 days, after which regular drinking water was returned. Mice were monitored daily at first, then weekly, and sacrificed at the indicated time points for tissue collection.

### ELISAs and qRT-PCR

*qRT-PCR*. Mucosal scrapings from the colons of mice were collected from DSS-treated mice. Briefly, entire colons (from proximal to distal end) were removed, cut longitudinally, and rinsed in ice-cold PBS to remove contents. Using the edge of a glass slide, the epithelial layer was separated from the muscularis propria and retained. For RNA isolation, 1 ml Trizol (ThermoFisher) was added to mucosal scrapings and immediately homogenized using a Pellet Pestle homogenizer (DWK Life Sciences, Millville, NJ). RNA isolation was performed according to manufacturer’s instructions. One ug total RNA was used for first strand cDNA synthesis of all samples Transcriptor High Fidelity cDNA kits (Sigma Aldrich) was used for first strand cDNA synthesis of all samples. For protein lysate, the epithelial layer was separated as described above, then immediately homogenized using the Pellet Pestle homogenizer in Cell Lysis Buffer (Cell Signaling). Lysate was processed according to manufacturer’s instructions. Protein concentrations of cellular lysates were determined using the Pierce BCA Protein Assay Kit from ThermoFisher.

*ELISAs*. Protein concentration of cellular lysates was first determined (Pierce BCA Protein Assay Kit #PI23227, ThermoFisher) before loading. 50 ug total protein of each sample were used in each well. The following ELISA kits were used: Hsp27 DuoSet ELISA (R&D Systems, DY1580) and ImmunoSet™ Hsp25 (rodent), ELISA development set (Enzo Life Sciences, ADI-960-075).

### Histology

Approximately 5 mm of the proximal and distal ends of the colon were cut from the tissue using a razor blade. Tissues were preserved overnight in either formalin, or Carnoy’s fixative (60% methanol, 30% chloroform, 10% glacial acetic acid) to preserve the mucus layer. The following morning, tissues were transferred to 70% ethanol until paraffin embedding. 5 um thick sections were cut. H&E staining was performed by the Human Tissue Resource Center at the University of Chicago.

For swiss rolls of SPF and GF colons, colon tissues were removed from mice as described above. Colons were cut longitudinally, pinned out flat on a bed of paraffin, and fixed with formalin for after 20 hours. While holding the proximal end of the colon with forceps, the tissue was rolled with up and held in place with a pin. Tissue was embedded in paraffin by The University of Chicago Human Tissue Resource Center. Sections were cut at 5 um thickness. To detect Hsp25 protein expression, we performed immunohistochemistry on IHC of Hsp25 antibody (Hsp25 ab, ImmunoSET, Hsp25 capture ab, cat #80-1942, ENZO) Dako IHC buffers, then counterstained with hematoxylin.

Histology scoring criteria: 0, no inflammation; 1, modest numbers of infiltrating cells in the lamina propria; 2, infiltration of mononuclear cells leading to separation of crypts and mild mucosal hyperplasia; 3, massive infiltration with inflammatory cells accompanied by disrupted mucosal architecture, loss of goblet cells, and marked mucosal hyperplasia; 4, presence of features in 1-3 plus crypt abscesses or ulceration. (Hu et al. Inflammation-induced, 3UTR-dependent translational inhibition of Hsp70 mRNA impairs intestinal homeostasis. Am J Physiol Gastrointest Liver Physiol 296: G1003–G1011, 2009.)

### IEC isolation and colonoid generation for *in vitro* stimulation assays

Mice were euthanized by CO_2_ asphyxiation. Entire colons were removed from mice and cut longitudinally. Strips of colon tissue were rinsed in ice-cold PBS to remove contents, then cut into small pieces. Colonic tissue was then washed several more times through a serological pipet until clean. The tissue pieces were incubated in 25 mM EDTA/PBS for 1 hour at 4°C on a rocking platform. Subsequently, the EDTA solution was aspirated and replaced with Advanced DMEM/F12 (ADF; ThermoFisher). Intestinal crypts were eluted by vigorous pipetting up and down. Remaining tissue pieces were filtered out using a 70 um cell strainer. Crypts were then centrifuged at 400xg for 15 min in a ThermoFisher Sorvall 16RT centrifuge at 4°C, then washed an additional time. The cell pellet was divided and resuspended into the respective wells for stimulation.

For colonoid generation, we used a similar procedure as described by Sato, et al. 2011 Gastroenterology, with a few modifications. Colons were removed from the mice as described above. The tissue pieces were incubated in 2.5 mM EDTA/PBS for 1 hour at 4°C on a rocking platform. Then, EDTA/PBS was aspirated and replaced with 2 ml (per colon) pre-warmed TryPLE Express (ThermoFisher) and incubated in a 37°C water bath. After 15 minutes, the digestion was arrested with the addition of ADF media. Intestinal crypts were eluted by vigorous pipetting up and down. Remaining tissue pieces were filtered out using a 70 um cell strainer. Crypts were then centrifuged at 400xg for 15 min in a ThermoFisher Sorvall 16RT centrifuge at 4°C. After washing, the cell pellet was resuspended in a 1:2 mixture of colonoid medium:Matrigel (Corning). Mouse colonoid medium contains 50% ADF, 50% L-WRN (ATCC), 1X N2 supplement (ThermoFisher), 1X B27 supplement (ThermoFisher), EGF (50 ng/ml, Peprotech), R-spondin-1 (50 ng/ml, Peprotech), Jagged-1 peptide (1 uM, Genscript), Y-27632 (10 uM, Selleck Chemicals), 1 mM HEPES (ThermoFisher), 1X Penicillin/Streptomycin (ThermoFisher), 1X L-glutamine (ThermoFisher). Colonoid media was changed every 2-3 days. Colonoids were cultured over several weeks with periodic splitting/expansion until enough cells were generated for the experiment. No colonoids with passage number higher than 10 were used.

For cytokine stimulation, recombinant murine IL-6 or IL-22 were purchased from Peprotech. Cells were stimulated at 37ºC/5% CO_2_ in 12-well plates for the indicated times with either IL-6 (100 ng/ml) or IL-22 (30 ng/ml) in ADF media.

### Western blotting, Immunoprecipitation, Silver staining

Antibodies used for western blots were: pSTAT3 Y705 (Cell Signaling #9145), pSTAT3 S727 (Cell Signaling #9134), STAT3 (Cell Signaling #9139 or #4904), YAP (Cell Signaling #4912), and Non-phospho (Active) YAP S127 (Cell Signaling #29495).

Protein concentration of all cellular lysates were first determined using the Pierce BCA Protein Assay Kit #PI23227 (ThermoFisher) prior to SDS-PAGE. For immunoprecipitation, lysates were prepared using the Pierce Classic Magnetic IP and CO IP kit #88804 (ThermoFisher). Anti-pSTAT3 Y705 (Cell Signaling #9145) and Hsp25 (Abcam #ab202846) were used to detect protein binding partners of pSTAT3 Y705 and Hsp25, respectively. To minimize interference of the IgG signal in the immunoprecipitation, we used Tidyblot (Bio-Rad #STAR209P). For silver stain gels, Silver Stain Plus Kit (Bio-Rad #1610449) was used.

## Acknowledgments

We thank The University of Chicago Human Tissue Resource Center (RRID:SCR_019199), especially Christy Schmehl and Xin Jiang, for their assistance with histological processing and staining.

## Supplemental Tables and Figure Titles and legends

**Figure S1.**
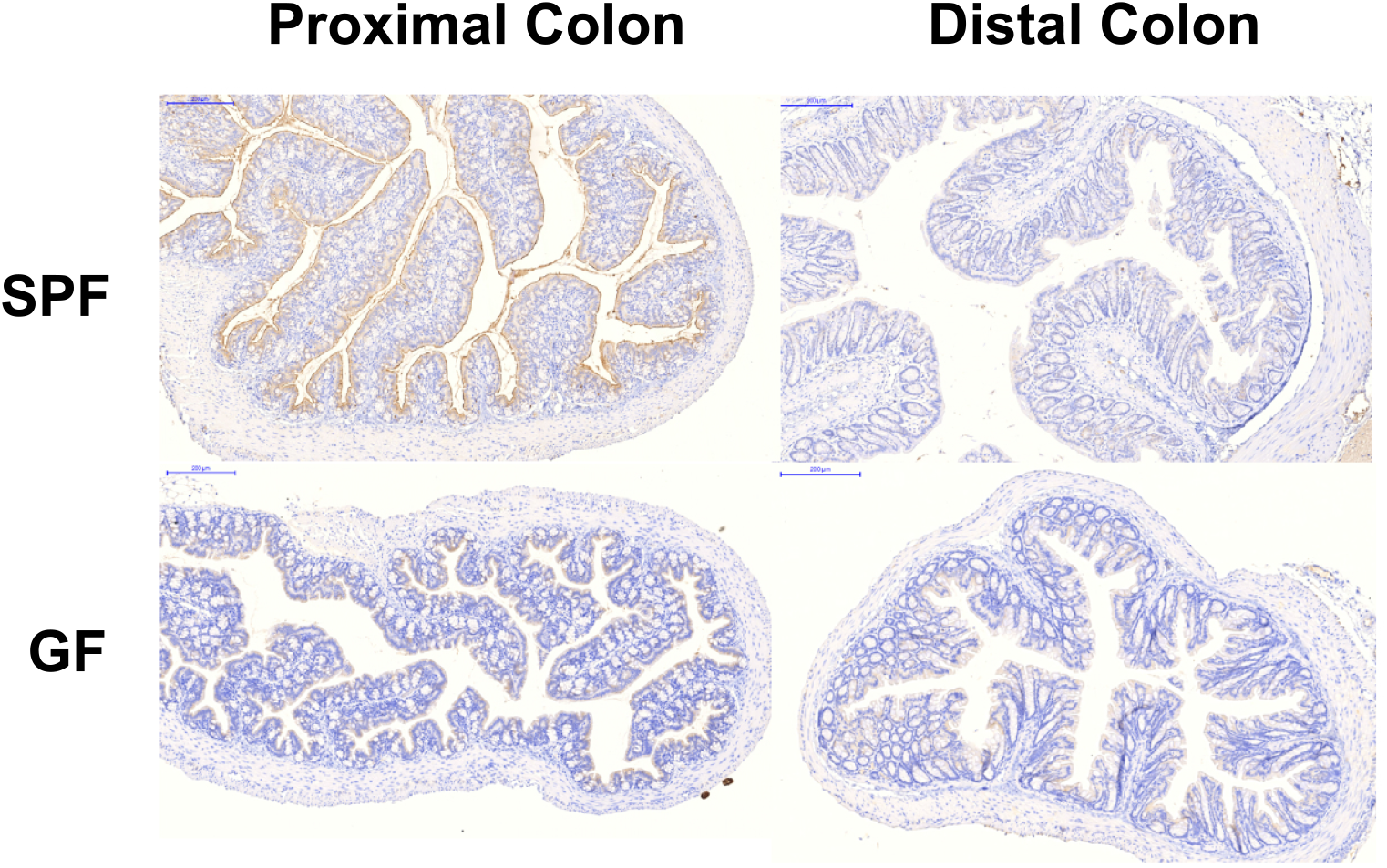
Hsp25 expression is regulated by microbes. Formalin-fixed, paraffin embedded proximal and distal colons from wild-type SPF and GF mice were subjected to staining with anti-pHSP25 S86.

**Figure S2.**
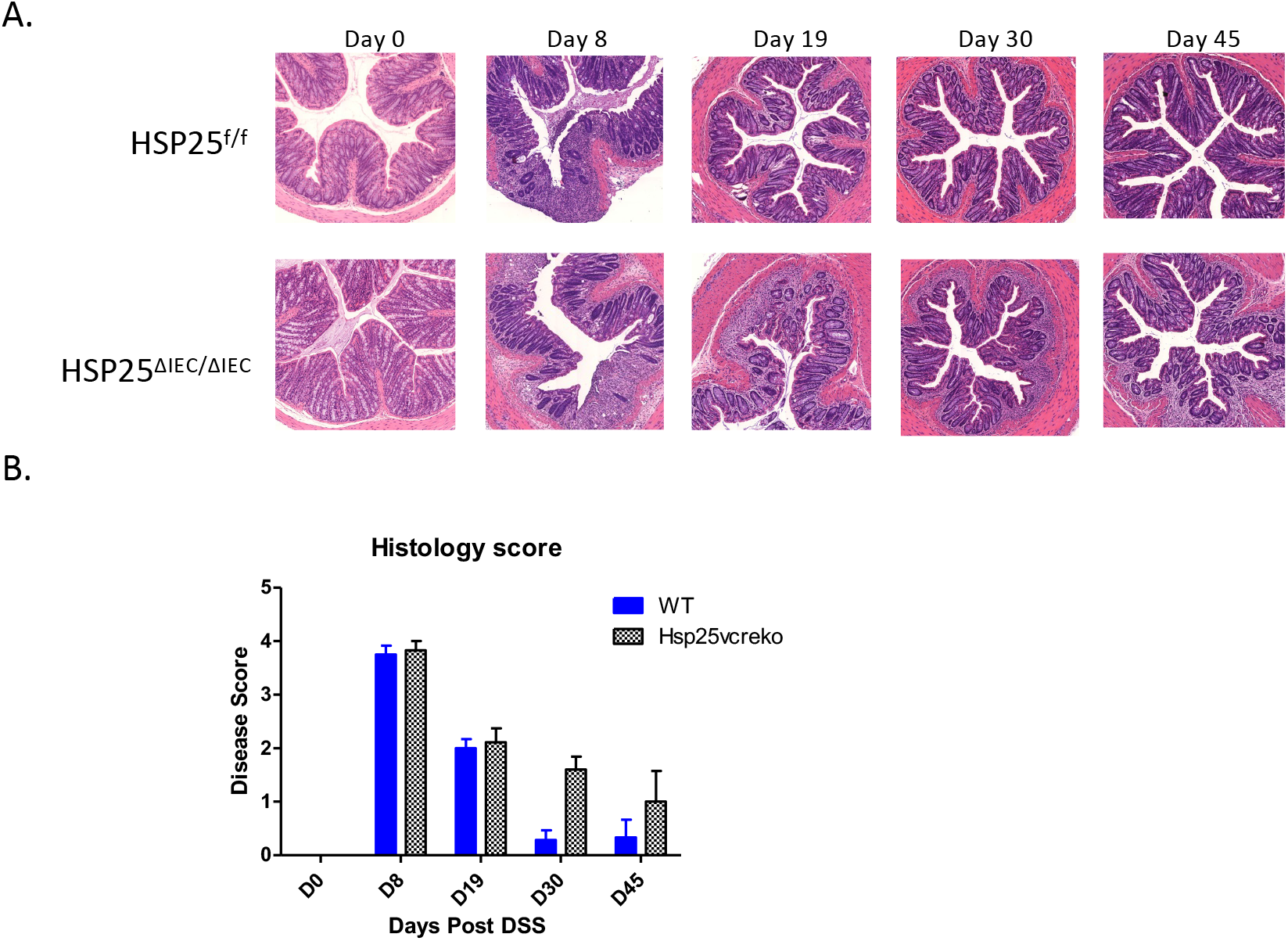
Hsp25^DIEC^ exhibit a similar phenotype as Hsp25 KO when treated with DSS. (A) Colonic mucosal histology of Hsp25f/f and Hsp25^DIEC^ mice treated with 2.5% DSS. (B) Histology scores of Hsp25f/f and Hsp25^DIEC^ mice treated with 2.5% DSS.

**Figure S3.**
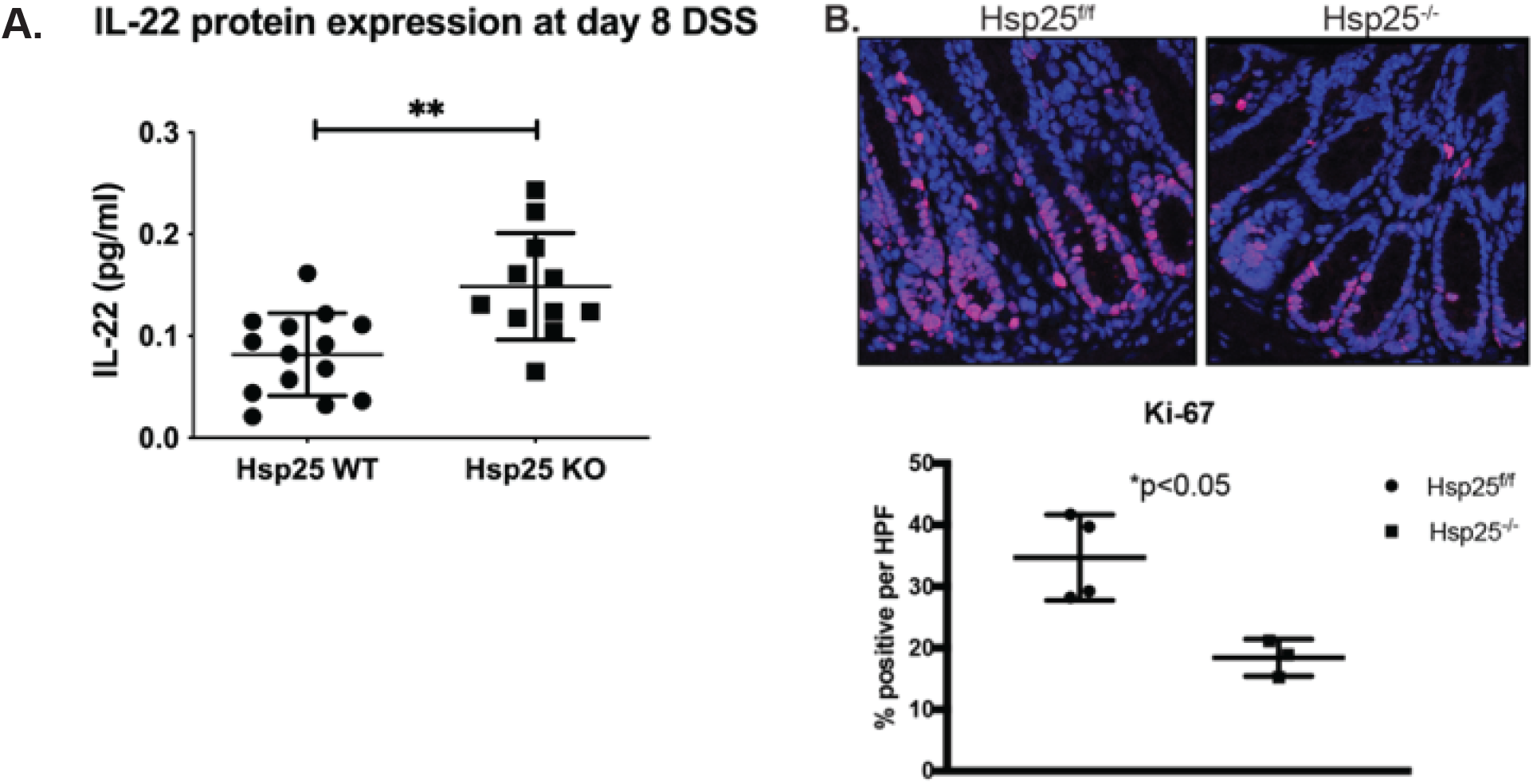
Increased IL-22 protein expression, but reduced Ki-67 staining in Hsp25 KO mice. (A) IL-22 protein levels from colonic mucosal scrapings of day 8 DSS-treated Hsp25 WT and Hsp25 KO mice as measured by ELISA. (B) Representative Ki-67 staining from day 19 DSS-treated Hsp25 WT and Hsp25 KO mice.

**Figure S4.**
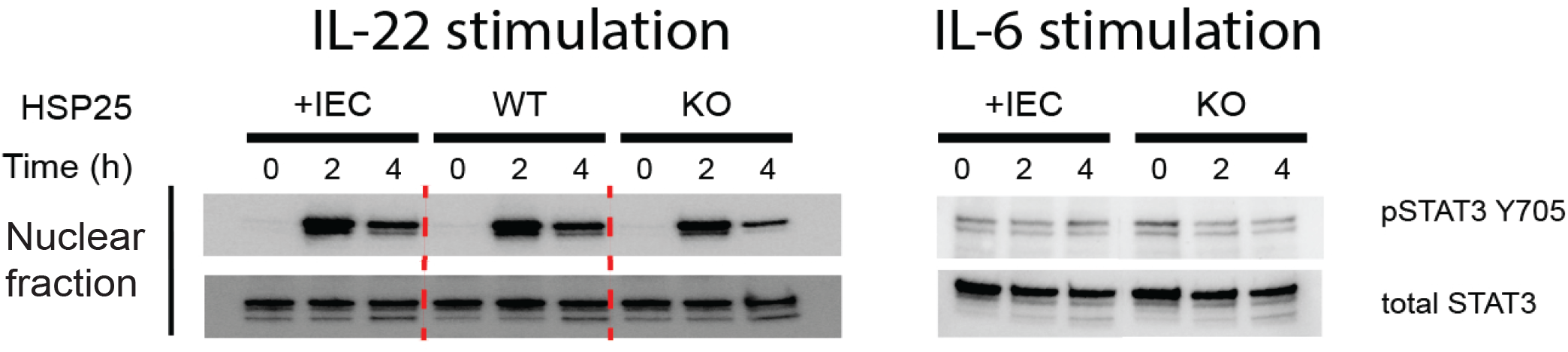
IL-22 and IL-6 stimulation of colonic IEC result in different phosphorylation activities of STAT3 at Y705. Colonic IECs from Hsp25^+IEC^, Hsp25 WT, and Hsp25 KO were stimulated with IL-22 (30 ng/ml) and IL-6 (100 ng/ml) for the indicated times. Cells were then collected and lysed into nuclear and cytosolic fractions. Nuclear pSTAT3 Y705 and total STAT3 levels were evaluated by Western blot.

